# Peripheral blood T-cell deficiency and hyperinflammatory monocyte responses associate with MAC lung disease

**DOI:** 10.1101/2022.03.25.485768

**Authors:** Cecilia S. Lindestam Arlehamn, Basilin Benson, Rebecca Kuan, Kimberley A. Dill-McFarland, Glenna J. Peterson, Sinu Paul, Felicia K. Nguyen, Robert H. Gilman, Mayuko Saito, Randy Taplitz, Matthew Arentz, Christopher H. Goss, Moira L. Aitken, David J. Horne, Javeed A. Shah, Alessandro Sette, Thomas R. Hawn

## Abstract

**Rationale:** Although nontuberculous mycobacterial (NTM) disease is a growing problem, available treatments are suboptimal and diagnostic tools are inadequate. Immunological mechanisms of susceptibility to NTM disease are poorly understood.

**Objective:** To understand NTM pathogenesis, we evaluated innate and antigen-specific adaptive immune responses to *Mycobacterium avium* complex (MAC) in individuals with MAC lung disease (MACDZ).

**Methods:** We synthesized 15mer MAC-, NTM-, or MAC/Mtb-specific peptides and stimulated peripheral blood mononuclear cells (PBMC) with pools of these peptides. We measured T-cell responses by cytokine production, expression of surface markers, and analysis of global gene expression in 27 MACDZ individuals and 32 healthy controls. We also analyzed global gene expression in Mav-infected and uninfected peripheral blood monocytes from 17 MACDZ and 17 healthy controls.

**Measurements and Main Results:** We were unable to detect T-cell responses against the peptide libraries or Mav lysate that has increased reactivity in MACDZ subjects compared to controls. T-cell responses to non-mycobacteria derived antigens were preserved.

MACDZ individuals had a lower frequency of Th1 and Th1* T-cell populations. By contrast, global gene expression analysis demonstrated upregulation of proinflammatory pathways in uninfected and Mav-infected monocytes derived from MACDZ subjects compared to controls.

**Conclusions:** Peripheral blood T-cell responses to Mycobacterial antigens and the frequency of Th1 and Th1* cell populations are diminished in individuals with MAC disease. In contrast, MACDZ subjects had hyperinflammatory monocyte responses. Together, these data suggest a novel immunologic defect which underlies MAC pathogenesis and includes concurrent innate and adaptive dysregulation.

## Introduction

Nontuberculous mycobacteria (NTM) are commonly encountered in the environment (1–3). Despite widespread exposure to NTMs, few humans develop disease. Risk factors for NTM lung disease include cystic fibrosis, structural lung disease, and a syndrome in women with higher rates of scoliosis, pectus excavatum, and a low body mass index (4, 5). Disseminated NTM infection is associated with Mendelian susceptibility to mycobacterial disease (MSMD; OMIM#209950), a rare pediatric disease caused by inborn errors of IFN*γ* immunity (6), suggesting a role for IFN*γ*. Although previous studies suggest that adults with MAC disease have low levels of IFN*γ*-production, the mechanisms underlying this cellular defect are not known (7–9) and the majority of NTM infections occur without identification of a genetic or immune defect.

Species in the *Mycobacteria* genus, which includes NTMs, the Bacille Calmette-Guerin (BCG) vaccine, and *Mycobacterium tuberculosis* (*Mtb*), share many similarities, including a lipid-rich cell wall and conserved proteins (10). Previous studies suggest that prior NTM exposure and sensitization could provide a protective immune response against tuberculosis (TB) (i.e., heterologous mycobacterial immunity) and/or impair BCG-induced vaccine protection (11–14). Animal models support that NTM infection is protective against TB disease and vice versa (14, 15). However, these previous studies were focused on a limited number of immune responses (IFNγ-production or delayed type hypersensitivity in skin), employed whole cell reagents rather than single proteins or peptides, and/or were underpowered. With ongoing vaccine developing efforts for TB, understanding the role of NTM exposure and heterologous immune responses may be critical for success (11).

Despite geographic variation of NTM, the *Mycobacterium avium* complex (MAC) including *M. avium* and *M. intracellulare,* are the most common NTM in all global regions (16), and the main drivers of the increasing incidence of NTM infection (17–19). Diagnosis of MAC lung disease is often challenging due to difficulties in collecting sputum and differentiating between MAC colonization and lung disease (20). Efforts to improve MAC diagnostics included skin tests (21, 22) and serodiagnostics (23–25). However, immunologic tests which accurately diagnose MAC, predict disease progression, or assess the success of treatment are not currently available. Detection of MAC antigen-specific T-cell responses could lead to an assay to identify MAC exposure, infection, and/or disease, and to test concepts of heterologous immunity.

In this study, we compared innate and adaptive immune responses to *M. avium* (Mav) in individuals with and without MAC lung disease (MACDZ). We hypothesized that Mav-specific immune responses are associated with susceptibility to MAC lung disease. We further tested whether it was possible to define T-cell epitopes associated with MAC-specific immune responses.

## Methods

### Study Participants and Ethics Statement

Approval for study protocols was obtained from the institutional review boards at the University of Washington School of Medicine and La Jolla Institute for Immunology. All participants provided written informed consent prior to participation in the study. We enrolled subjects in Seattle who had a history of MAC isolated from a sputum sample. Among the MAC subjects, the majority met ATS criteria for MAC lung disease (20) with a history of both pulmonary symptoms at the time of diagnosis and abnormalities on chest radiography (Table 1). PBMCs were obtained months to years after the completion of treatment at a time without symptoms. We designed two study groups for adaptive (T-cell response) and innate (Monocyte Mav infection) immune profiling (Table 1). For innate profiling, we also enrolled local controls in Seattle who were self-described as healthy without history of recurrent or serious infections. For adaptive profiling, we enrolled healthy controls at the University of California, San Diego Anti-Viral Research Center (San Diego, USA, n=28), and at the Universidad Peruana Cayetano Heredia (Lima, Peru, n=4), both with (IFN*γ*-release assay (IGRA)+HC) and without (IGRA-HC) latent tuberculosis infection. Mtb infection status was confirmed by a positive IGRA (QuantiFERON-TB Gold In-Tube, Cellestis, or T-SPOT.TB, Oxford Immunotec) and the absence of symptoms consistent with TB or other clinical and radiographic signs of active TB.

**Table 1:** Demographic and clinical characteristics of cohorts.

|  | **Monocyte *M. avium***<br>**Infection** |  | **MAC/NTM T-cell Response** |  |  |
| --- | --- | --- | --- | --- | --- |
|  | **HC**<br>**(N=17)<sup>1</sup>** | **MACDZ**<br>**(N=17)** | **IGRA- HC**<br>**(N=19)** | **IGRA+ HC**<br>**(N=13)** | **MACDZ**<br>**(N=27)** |
| **Gender** |  |  |  |  |  |
| F (%) | 12 (70.6) | 14 (82.4) | 9 (47.4) | 3 (23.1) | 16 (59) |
| M (%) | 5 (29.4) | 3 (17.6) | 10 (52.6) | 10 (76.9) | 11 (41) |
| **Cystic Fibrosis** |  |  |  |  |  |
| Absent (%) | 17 (100) | 11 (64.7) | N/A | N/A | 12 (44) |
| Present (%) | 0 (0) | 6 (35.3) | N/A | N/A | 15 (56) |
| **Age** |  |  |  |  |  |
| Mean (SD)<sup>2</sup> | 53.4 (7.98) | 58.2 (15.0) | 39.7 (12.1) | 44.2 (14.5) | 49.1 (20.7) |
| Median | 55.0 | 62.0 | 41.0 | 44.0 | 45.0 |
| [Min<sup>3</sup>, Max<sup>4</sup>] | [42.0, 65.0] | [35.0, 84.0] | [22, 60] | [22, 62] | [26, 82] |
| **Ethnicity** |  |  |  |  |  |
| Asian (%) | 2 (11.8) | 0 (0) | 3 (15.8) | 0 (0) | 0 (0) |
| Black (%) | 1 (5.9) | 0 (0) | 1 (5.3) | 5 (38.5) | 1 (3.7) |
| Pacific Islander (%) | 0 (0) | 1 (5.9) | 0 (0) | 0 (0) | 0 (0) |
| White (%) | 14 (82.4) | 16 (94.1) | 10 (52.6) | 1 (7.7) | 25 (92.6) |
| Mixed (%) | 0 (0) | 0 (0) | 5 (26.3) | 6 (46.2) | 0 (0) |
| Not reported (%) | 0 (0) | 0 (0) | 0 (0) | 1 (7.7) | 1 (3.7) |
| **Diabetes Mellitus** |  |  |  |  |  |
| Absent (%) | 17 (100) | 15 (88.2) | N/A | N/A | 21 (77.8) |
| Present (%) | 0 (0) | 2 (11.8) | N/A | N/A | 5 (18.5) |
| Not reported (%) | 0 (0) | 0 (0) | 19 (100) | 13 (100) | 1 (3.7) |
| **Malignancy** |  |  |  |  |  |
| Absent (%) | 17 (100) | 16 (94.1) | N/A | N/A | 27 (100) |
| Present (%) | 0 (0) | 1 (5.9) | N/A | N/A | 0 (0) |
| **Bronchiectasis** |  |  |  |  |  |
| Absent (%) | N/A | 5 (33.3) | N/A | N/A | 4 (14.8) |
| Present (%) | N/A | 10 (66.6) | N/A | N/A | 22 (81.5) |
| **PLHIV<sup>5</sup>** |  |  |  |  |  |
| Absent (%) | 17 (100) | 16 (94.1) | 19 (100) | 13 (100) | 27 (100) |
| Present (%) | 0 (0) | 1 (5.9) | 0 (0) | 0 (0) | 0 (0) |
| **Chest Radiographic Abnormality<sup>6</sup>** |  |  |  |  |  |
| Absent (%) | N/A | 0 (0) | N/A | N/A | 1 (3.7) |
| Present (%) | N/A | 14 (82.3)<sup>7</sup> | N/A | N/A | 26 (96.3) |

<sup>1</sup> Numbers not adding to 100% are due to missing data. One individual with MAC overlapped between the T-cell and monocyte studies. <sup>2</sup> SD, Standard Deviation. <sup>3</sup> Min, Minimum. <sup>4</sup> Max, Maximum. <sup>5</sup> HIV, Person Living with Human Immunodeficiency Virus. <sup>6</sup> Abnormalities include ground glass opacities, nodule, infiltrate, bronchiectasis. <sup>7</sup> Three individuals without available CXR data.

### Peptides

To discover the epitopes targeted by MAC-specific T-cell responses in MACDZ individuals, we constructed a candidate peptide library. A bioinformatic analysis was performed to define HLA class II binding 15-mer peptides from the genomes of different mycobacterial species. We identified protein sequences from NCBI and after removal of redundant sequences, had a total of 31,576 sequences from 7 representative species of MAC (Taxonomy id:120793), 11,340 from 7 representative species from the Mtb complex (Taxonomy ID: 77643) and an additional 286,183 from 57 other Mycobacteria species. Among all possible 15-mers, peptides that met the following criteria of three individual categories were selected: MAC-specific (only present in MAC strains), NTM-specific (not present in Mtb), and MAC/Mtb-specific (present in both MAC and Mtb). Next, class II binding prediction was performed using the 7-allele method (26), and peptides with a median percentile rank *≤*2 were selected. Any peptides from “hypothetical proteins” were excluded. This resulted in a peptide library of a total of 1,584 peptides: 628 MAC-specific, 516 NTM-specific, and 440 MAC/Mtb-specific (**Table S1**). Peptides were randomly divided into three peptide pools per category, resulting in nine pools.

Other pools included a previously described *Mtb*-derived pool (MTB300, (27)) which includes peptides found in NTM, a pool of EBV/CMV HLA class II-restricted epitopes (EBV/CMV-II; (28)) and one of Tetanus toxoid (TT; (29)) epitopes. Peptides were synthesized by A&A LLC (San Diego). Peptides were pooled into peptide pools, re-lyophilized and reconstituted at a concentration of 1mg/ml or 0.7mg/ml (MTB300).

The remainders of all experimental procedures are described in detail in the Supplemental Methods.

## Results

### MACDZ have infrequent MAC-antigen or mycobacteria-specific T-cell responses

To define T-cell responses against MAC antigens, we tested PBMCs from 10 MACDZ and 10 IGRA+HC (**Table 1**) with MAC-, NTM-, and MAC/Mtb-specific peptides, and MTB300 (27) which includes peptides found in NTMs, an EBV/CMV-II and a TT pool of epitopes as controls (Methods), as well as Mav and Mtb whole cell lysates.

The antigen-specific reactivity was assayed directly ex vivo using an IFN*γ*/IL-5/IL-17 Fluorospot assay (**Figure 1A**). Surprisingly little reactivity was detected against the pools in MACDZ individuals. As expected, the IGRA+HC also did not react to the 9 different peptide pools. The IGRA+HC had higher reactivity against Mtb lysate and MTB300 (as expected), but also a trend towards higher reactivity against Mav lysate. Both cohorts had similar reactivity against the non-mycobacteria derived peptide pools EBV/CMV and TT. The reactivity detected in both cohorts were primarily driven by IFN*γ*-specific responses, with barely any IL-5 or IL-17 detected (**Figure S2A**).

**Figure 1.**
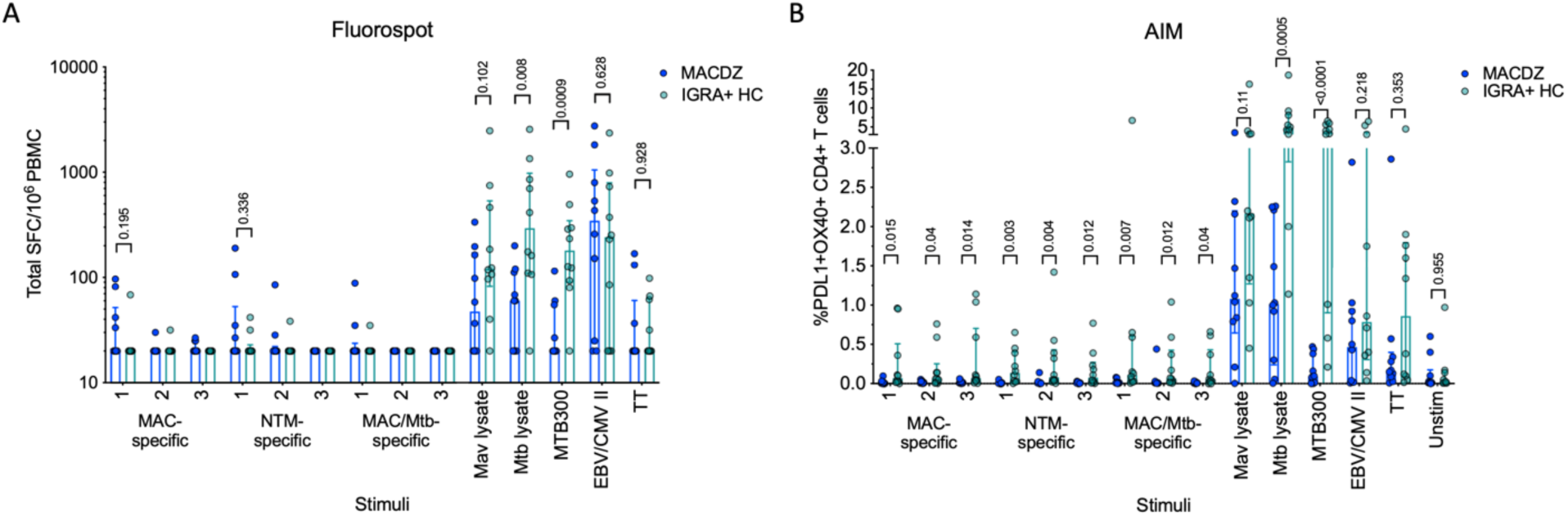
Individuals with MAC disease have infrequent Mav- or mycobacteria- specific responses. A) Total magnitude of response (sum of IFN*γ*, IL-5 and IL-17) against peptide pools, Mav and Mtb lysates as SFC per 10^6^ cultured PBMC as determined by Fluorospot. Each point represents one participant (MACDZ, n=10 in blue; IGRA+HC, n=10 in teal), median ± interquartile range is shown. Two-tailed Mann-Whitney test, comparisons without a p-value indicated were not significant. Limit of detection is 20 SFC. B) Frequency of PDL1+OX40+ CD4 T-cells in response to peptide pools, Mav and Mtb lysates. Each point represents one participant (MACDZ, n=10; IGRA+HC, n=10), median ± interquartile range is shown. Two-tailed Mann-Whitney test. CD4 T-cells were gated as CD3+CD4+CD8-CD19-CD14- in the live singlet gate of PBMC.

To determine whether reactivity in MACDZ was driven by a response other than IFN*γ*/IL-5/IL-17, we also used a cytokine-agnostic approach measuring Activation Induced Marker (AIM) upregulation following antigenic stimulation (**Figure 1B**). Upregulation of both OX40 and PDL1 has previously been used to measure Mtb-specific T-cell reactivity (30). Again, we found minimal reactivity against the 9 different peptide pools in both cohorts and the same hierarchy of responses against the controls (**Figure 1B**). IGRA+HC had a trend towards higher reactivity against Mav lysate irrespective of the activation markers investigated (**Figure S2B**). Finally, we measured stimulus-specific IFN*γ*, IL-2, TNF*α* and CD154 responses to determine whether MACDZ had a different polyfunctional response (**Figure S2C**), but as before, if anything, responses in the IGRA+HC were higher.

In conclusion, we were unable to define T-cell responses against the peptide library or increased reactivity compared to IGRA+HC against Mav lysate with multiple antigen-specific assays. Importantly, the lack of response did not translate more broadly to non-mycobacteria derived antigens.

### MACDZ have lower frequencies of specific PBMC cell subsets

To determine the cause for the lack of reactivity in MACDZ against Mav reagents, we determined basal frequencies of major PBMC subsets, e.g. measured without antigen stimulation in MACDZ (n=19) compared to IGRA-HC (n=18). We first analyzed the relative frequency of monocytes, NK cells, B cells, CD56-expressing T-cells, T-cells, CD4+ and CD8+ T-cells (**Figure 2A**). The frequency of monocytes was higher in MACDZ compared to IGRA-HC (p=0.036; Two-tailed Mann-Whitney test), and in contrast, the frequency of lymphocytes was lower (p=0.038). This difference was primarily driven by lower frequencies of CD8+ T-cells (p=0.0002), as all other cell subsets had similar frequencies (p>0.05).

**Figure 2.**
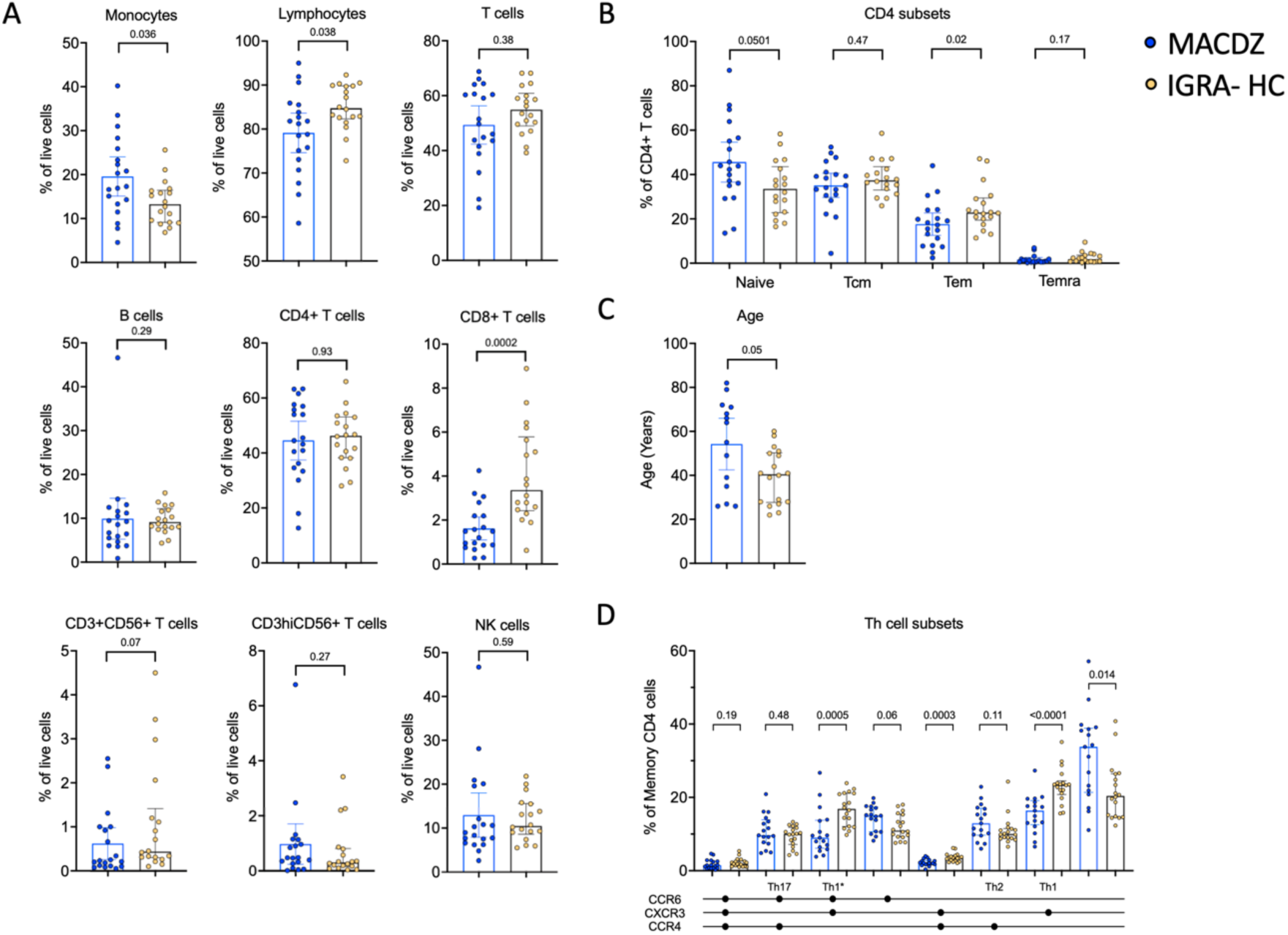
Individuals with MAC disease have lower frequencies of specific cell subsets. A) Frequency of major PBMC subsets in MACDZ (n=19, in blue) and IGRA-HC (n=18, in yellow). B) Frequency of CD4 memory populations based on CD45RA and CCR7 expression divided in naïve, effector memory (Tem), central memory (Tcm) and Temra populations. C) Age of the participants. D) Frequency of Th cell subsets based on CCR4, CCR6 and CXCR3 expression. A-D) Each point represents one participant, median ± interquartile range is shown. Two-tailed Mann-Whitney test.

We next analyzed the frequencies of CD4 and CD8 memory T-cell subsets (**Figure 2B, Figure S3A**). There was no significant difference between CD8 memory T-cell subsets (**Figure S3A**); however, for CD4 memory a significant lower level of Tem (CD45RA-CCR7-, p=0.02), and a higher level of naïve (CD45RA+CCR7+, p=0.05) T-cells was detected in MACDZ compared to IGRA-HC (**Figure 2B**). This was striking since MACDZ were older than IGRA-HC (p=0.05, **Figure 2C**) and we had hypothesized the opposite based on the general trend toward a shrinking naïve T-cell pool as a function of older age.

We also measured the frequency of T helper subsets based on the expression of CXCR3, CCR6, and CCR4. MACDZ had lower levels of Th1 (p<0.0001), Th1* (p=0.0005) and CXCR3+CCR6-CCR4+ (p=0.0003) memory CD4 T-cells, and conversely higher levels of CXCR3-CCR6-CCR4- (p=0.014) T-cells (**Figure 2D**). There were 9 individuals out of the 17 MACDZ tested with the Th subset markers that were affected by CF. There were no significant differences in Th1* and Th1 subsets in individuals with CF (**Figure S3B**). The lower frequency of Th1* and Th1 subsets, which are involved in *Mtb*-specific immune responses, is consistent with the lack of detectable Mycobacteria-specific T-cell responses in MACDZ.

### Transcriptional analysis of MACDZ reveals specific gene signatures

To further address MAC-specific immune responses, we performed RNAseq on PBMCs from the MACDZ (n=15), IGRA+HC (n=10) and IGRA-HC (n=15) cohort, after a 24 hours stimulation with Mav lysate, Mtb lysate, MTB300, and anti-CD3/CD28 as a positive control. The OX40/PDL1 AIM assay yielded results similar to those described above (**Figure S4A**). The MAC cohort was associated with the lowest number of OX40+PDL1+ CD4 T-cells in response to Mycobacteria-derived stimulation, but no difference in the response following anti-CD3/CD28 stimulation.

We performed differential gene expression analysis comparing MACDZ to IGRA+/-HC for unstimulated, MTB300-, Mav lysate-, and Mtb lysate-stimulated samples. Upon Mav-lysate and Mtb-lysate stimulation, we identified few differentially expressed genes (DEG) between the cohorts; 7 genes were upregulated in IGRA+/- individuals following Mav lysate stimulation and 41 genes were differentially expressed following Mtb lysate stimulation (FDR values <0.05; Benjamini Hochberg corrected Wald test, and log2 fold change >0.5 or <-0.5; **Table S2, Figure S4B**).

In contrast, gene signatures differentiating between MACDZ and IGRA+/-HC were found for both unstimulated and MTB300-stimulated samples. For unstimulated samples (**Figure 3A**), 124 genes were upregulated in MACDZ and 259 genes were upregulated in IGRA+/-HC. For MTB300- stimulated samples (**Figure 3B**), 123 genes were upregulated in MACDZ and 228 genes were upregulated in IGRA+/-HC (**Table S2**).

**Figure 3.**
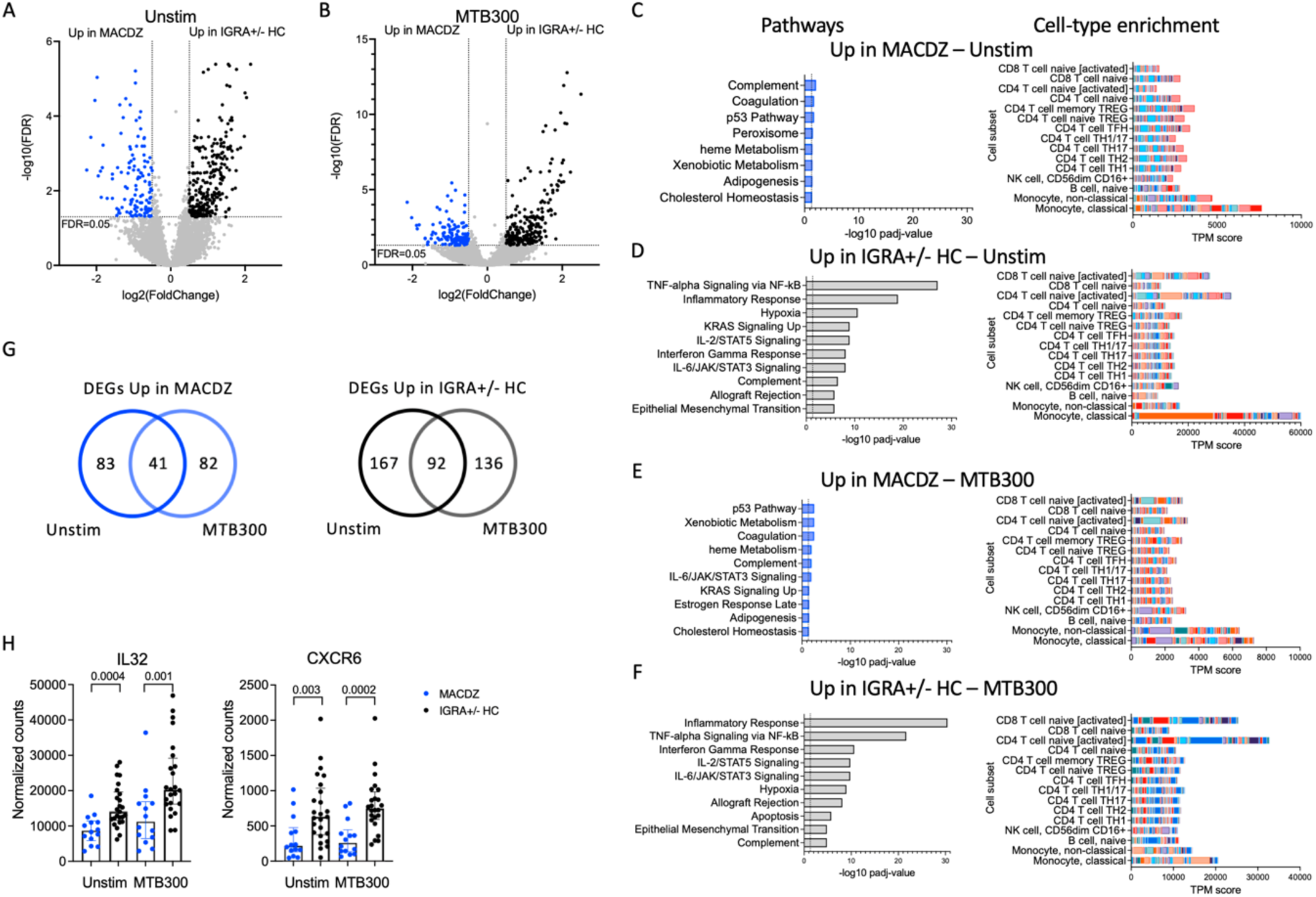
Specific gene signatures in individuals with MAC disease. A, B) Volcano plot showing differentially expressed genes in unstimulated samples (A) and MTB300 stimulated samples (B) comparing individuals with MAC disease (upregulated genes shown in blue) to IGRA+/-HC individuals (upregulated genes shown in black). FDR <0.05 and log2 fold change >0.5 or <-0.5, Benjamini Hochberg corrected DESeq2 Wald test. C-F) Pathway and cell-type enrichment (dice-database.org) for genes upregulated in unstimulated MAC samples (C), upregulated in unstimulated IGRA+/-HC samples (D), upregulated in MTB300 stimulated MACDZ samples (E), and upregulated in MTB300 stimulated IGRA+/-HC samples (F). Up to the ten most significant pathways are shown. G) Venn-diagrams showing overlap between DEGs for unstim and MTB300 stimulated MACDZ samples (left) and IGRA+/-HC samples (right). H) IL32 and CXCR6 gene expression in the unstimulated or MTB300 stimulated MACDZ and IGRA+/-HC samples. Each point represents one participant, median ± interquartile range is shown. Two-tailed Mann-Whitney test.

Hypergeometric mean pathway enrichment using Hallmark gene sets for the upregulated genes for each group showed similar pathways between unstimulated and MTB300-stimulated samples. Genes involved in heme metabolism, coagulation and complement were identified for MACDZ, although with adjusted p-values just above the cut-off of 0.05 (corresponding to -log10 1.3 in the figures). For IGRA+/-HC, we found significant enrichment for inflammatory response, interferon-gamma response and TNF-alpha signaling via NF-*κ*B (**Figure 3C-F**).

The similar pathways within each cohort in unstimulated vs. MTB300 stimulated samples was explained by a large overlap of DEGs (**Figure 3G, Table S2**). Using cellular deconvolution methods (through dice-database.org) for the upregulated genes in each cohort, we found an enrichment corresponding to classical and non-classical monocytes in MACDZ and, in contrast, an enrichment of activated CD4 and CD8 T-cells in IGRA+/-HC, which was, as expected, more pronounced following MTB300 stimulation (**Figure 3C-F**).

Several Th1*-related genes were upregulated in IGRA+/- individuals, 7 in unstimulated (p=0.11; overlap between upregulated genes here compared to the previously described Th1* signature (31)) and 9 in MTB300 stimulated samples (p=0.007), thus reflecting the differences observed in the phenotypic analysis described above. The IL-32 and CXCR6 expression was increased in IGRA+/-HC compared to MACDZ in both unstimulated and MTB300-stimulated samples (**Figure 3H;** remaining Th1* genes in Figure S4C). Overall, these results demonstrate a difference in MTB300-stimulated PBMC responses between MACDZ and the other groups, and link the cell subset frequency differences to the gene expression changes.

### Uninfected and Mav-infected monocytes in MACDZ have upregulated pro-inflammatory pathways

Together, these data suggested differences in monocyte frequency and monocyte response to MTB300 within PBMCs when comparing MACDZ and controls. We hypothesized that MACDZ have a myeloid cell functional alteration which underlies the deficient T-cell responses. We enriched CD14+ monocytes from PBMC (MACDZ, N=17; N=14 with MAC lung disease, 3 without CXR data available; or HC, N=17, **Table 1**), infected with *M. avium* (Mav, UCLA strain 104) or media for 6 hours, and obtained RNAseq transcriptional profiles. Using an interaction model that included clinical phenotype (MACDZ versus HC) and Mav infection (Mav versus media), we identified 227 DEGs; 138 were significant for MACDZ only (Mav independent), 87 were significant for MACDZ and Mav infection, and 2 were significant for the interaction term (together being Mav-dependent; FDR <0.05, **Figure 4A, Table S3**).

**Figure 4.**
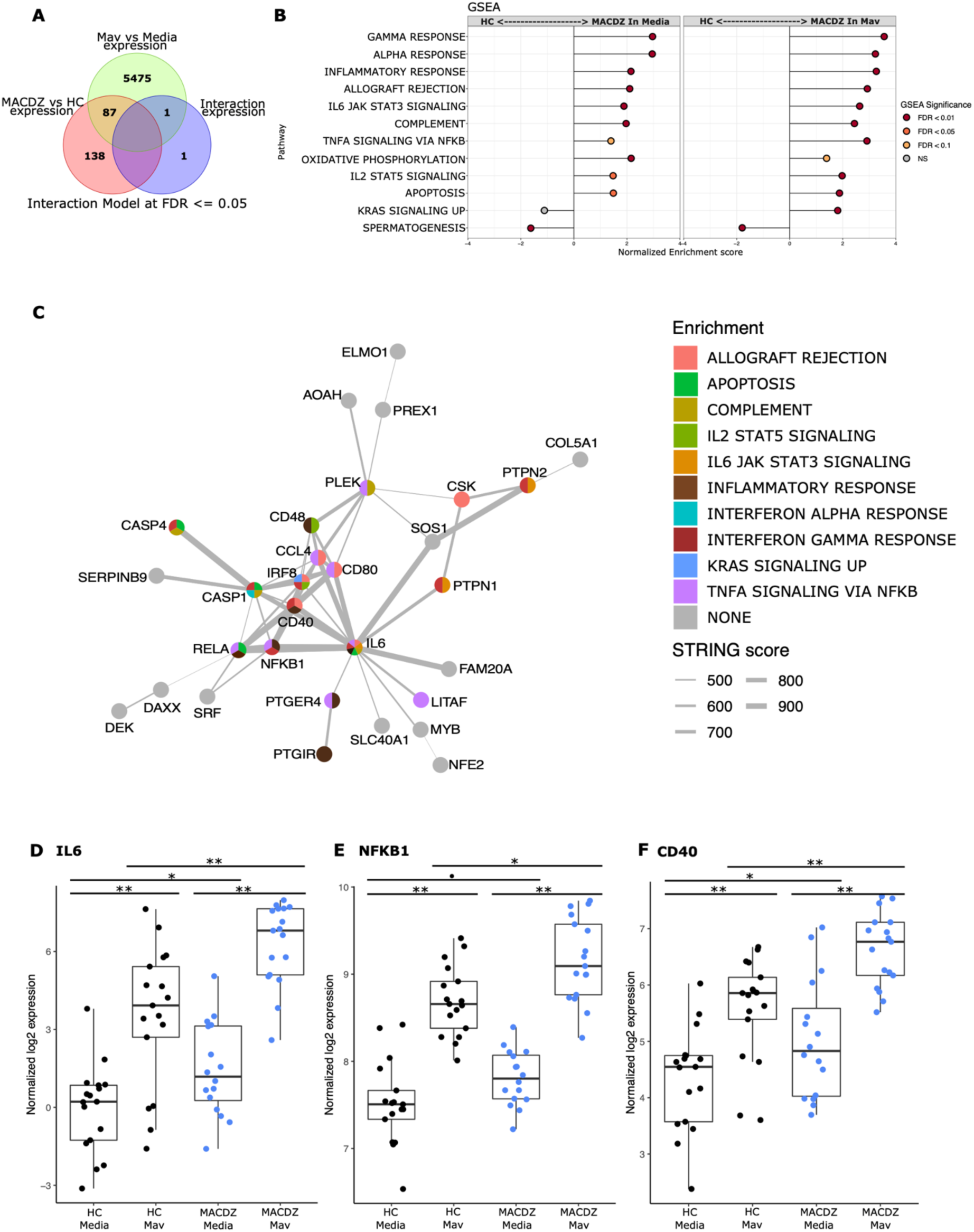
Monocyte response to *Mycobacterium avium* infection in MACDZ versus HC subjects. RNASeq transcriptional profiles were measured from monocytes isolated from MACDZ subjects (N=17) or healthy controls (N=17) with a media only condition or after infection with *M. avium* (MOI=5) for 6 hours. A) Expression profiles were compared between MACDZ and HC subjects with and without Mav infection using a linear model with an interaction term. Venn diagram depicts 227 differentially expressed genes (DEGs) distinguishing MACDZ vs HC subjects included 138 Mav-independent DEG significant for MACDZ alone, 87 for both MACDZ and Mav, and 2 for the interaction term, together defined as Mav-dependent DEG (FDR <0.05). B) Gene set enrichment analysis (GSEA) with Hallmark terms comparing MACDZ vs HC subjects within media or Mav infection conditions. C) STRING network analysis of 89 Mav-dependent DEGs (87 Mav plus 2 from the interaction term) which differentiate MACDZ vs HC subjects. Circles depict genes and are colored by membership in Hallmark mSigDB gene sets. Grey lines depict annotated connection between 2 genes in the STRING database with line thickness proportionate to the score. D-F) Boxplots depicting voom normalized log2 mRNA expression values in MACDZ versus HC subjects with and without Mav infection. FDR values depict comparison of MACDZ vs HC media expression, MACDZ vs HC Mav expression, and Mav vs media for MACDZ or HC subjects (FDR*≤*0.1 shown by •, FDR*≤*0.05 shown by *, and FDR*≤*0.01 shown by **). Median and interquartile range depicted.

To discover gene signatures that distinguished MACDZ vs. HC monocyte populations, we employed gene set enrichment analysis (GSEA) with the entire dataset (32) using the molecular signatures database (MSigDB) ‘Hallmark’ curated gene sets (33). We identified multiple gene signatures that differentiated MACDZ and HC monocytes, including an enrichment of inflammatory and signaling genes upregulated in MACDZ compared to HC, in both unstimulated and Mav infected monocytes (**Figure 4B, Table S4**). The most significantly enriched gene sets included Inflammatory Response, Interferon-Gamma Response, TNFA Signaling via NFKB, and IL6 JAK STAT3 Signaling (FDR < 0.01 for media, Mav, or both).

We also used hypergeometric mean pathway enrichment for the 138 Mav-independent and 89 Mav-dependent DEGs. Only Mav-dependent DEGs showed significant enrichment; this included the same pathways as GSEA, namely Inflammatory Response, Interferon-Gamma Response, and TNFA signaling via NFKB (**Table S5**). Analysis of the most significantly enriched KEGG pathways provided additional details on more specific pathways including Chemokine_Signaling_Pathway, Cytosolic_DNA_Sensing_Pathway, Nod_Like_Receptor Signaling_Pathway, and Toll-Like_receptor_ Signaling_Pathway (FDR < 0.001, Table S5). Taken together, these data suggest that monocyte transcriptional profiles of MACDZ are enriched for pro-inflammatory pathways compared to HC in both media and Mav infection conditions.

To further explore biologic pathways connected to these DEGs, we used network analysis (string-db.org) to connect the DEGs. For Mav-independent DEGs, we found a cluster (N=28 genes) of highly interconnected DEGs and 61 DEGs that had three or fewer connections (**Figure S7**). For the Mav-dependent DEGs, we found a different highly interconnected cluster (N=31 genes) that was centered on IL-6 and contained transcription factors (REL, NF-kB, IRF8, NFE2, and SRF), cytoplasmic signaling molecules (CASP1, CASP4), co-stimulatory molecules (CD40, CD48, CD80), cytokines (IL6), and chemokines (CCL4, CCL8) (**Figure 5C**). For each of these genes, expression values were higher in MACDZ compared to HC for the media and/or Mav condition (**Figure 5D-F**, **Figure S8**). Overall, our results suggest that MACDZ have a higher pro-inflammatory expression profile in monocytes in both unstimulated and Mav-infected conditions.

## Discussion

We present a comprehensive characterization of the innate and adaptive immune responses in MACDZ individuals. First, we found attenuated T-cell responses across multiple cellular subsets, including diminished responses of the Th1* subset that is important for antimycobacterial immunity (34, 35). Second, we were unable to detect T-cell responses against MAC-specific peptides or increased reactivity compared to IGRA+HC against Mav lysate in multiple antigen-specific assays. Third, transcriptional analysis of Mav-infected blood monocytes demonstrated enhanced innate immune activation in MACDZ compared to controls. To our knowledge, these are the first concurrent observations of both hyperinflammatory innate and hypoinflammatory adaptive profiles in MACDZ subjects, which provide a new conceptual framework for understanding MAC immunopathogenesis.

Previous studies demonstrated that MAC disease occurs in individuals with T-cell defects, such as those with AIDS, rare genetic or autoimmune diseases which comprise IFN*γ*-dependent immune responses (6). Furthermore, the lack of MAC-specific IFN*γ* response is in concordance with previous studies where low IFN*γ* was detected following lysate or antigen-mixture stimulation in individuals with pulmonary NTM/MAC without previously identified immune defects (7–9, 36). We extend these studies with the identification of specific T-cell subsets underlying the defect, discovery of concurrent hyperinflammatory innate responses, and use of peptide reagents with well-defined specificity. We were not able to define MAC- or NTM-specific T-cell epitopes in this study in individuals with known MAC exposure and lung disease. This could support a model whereby MACDZ do not contain immunodominant T-cell epitopes which are species-specific. However, previous work from our group and others have defined NTM-specific T-cell reactivity in healthy individuals without Mtb infection, with an assumed NTM exposure (35, 37). Here, we attempted to define these responses at a MAC-species peptide-specific level in subjects with known MAC disease and presumed T-cell sensitization. Despite our use of unique sets of MAC-, NTM-, and MAC/Mtb-specific peptides, similar to previous studies predicted for promiscuous HLA class II binding, we did not detect any T-cell responses that were enriched in MACDZ. Together, these data and prior studies suggest that the lack of response is due to a host response that is specific to those with documented MAC disease.

Our results point towards a role for Th1* T-cells in MACDZ. The Th1* subset contains the majority of *Mtb*- and NTM-specific T-cells (31, 35, 38), and a lower frequency of this subset could lead to a lack of Mycobacteria-specific responses. This hypothesis is strengthened by the previous observation that IGRA+ individuals have an increased frequency of the Th1* subset compared to IGRA- controls (31) and that this subset mediates BCG-induced CD4 T-cell responses and is increased following vaccination (39). The underlying cause for the lower frequency of Th1* and Th1 subsets in MACDZ remains to be determined, but could also predispose these individuals to this unusual infection. Additionally, excessive antigenic exposure from persistent MAC infection and inflammation could dampen T-cell function due to terminal differentiation, or “exhaustion”. In *Mtb* infection, increased frequency of terminally-differentiated PD1-KLRG1+CD4+ T-cells is associated with increased bacterial burden (40). The combination of ongoing infection in an excess of proinflammatory innate immune responses provides evidence for T-cell dysfunction. Another contributing factor includes anatomic barriers. Many MACDZ individuals, those with ciliary dysfunction, cystic fibrosis, or preexisting bronchiectasis (11, 41), have anatomic lung defects that separate bacilli from adaptive immune surveillance. Our data suggest that separation of bacilli from T-cells by thick mucus or absent lung tissue may prevent T-cell activation. These observations are consistent with effector memory T-cells not replenishing over time without ongoing antigenic stimulation (42).

Surprisingly, we also observed that blood monocyte pro-inflammatory responses, were enhanced in MACDZ, despite the lack of effective T-cell help. These data are consistent with several immunologic models. First, enhanced pro-inflammatory innate responses could stimulate persistent T-cell activation via cytokines and contribute to T-cell exhaustion and terminal differentiation. Alternatively, these enhanced innate responses could promote epigenetic changes in myeloid cells which result in trained immunity features (43). In murine trained immunity studies, BCG infected bone marrow and influenced macrophage differentiation and function to induce increased innate responses to a broad array of pathogens (43, 44). Although there is currently no evidence for trained immunity in human MAC disease, our studies suggest that MAC disease is associated with decreased T-cell responses, which permits Mav replication leading to alterations of macrophage responses during chronic infection. Alternatively, MACDZ myeloid cells may be genetically programmed with accentuated inflammatory responses which drive susceptibility to MAC disease. Recent genetic studies support this possibility, although the genes underlying MAC susceptibility remain poorly understood (45–47). Further studies will be needed to assess the impact of both macrophage hyperreactivity and relative T-cell deficiency in Mav pathogenesis.

While no diagnostic can fully supplant microbiologic testing, non-sputum-based biomarkers for patient screening, to predict progressive lung disease, and response to treatment would greatly advance the clinical care of patients with NTM disease (48, 49). In addition to our findings that MACDZ had a higher pro-inflammatory expression profile in monocytes, Cowman identified over 200 transcripts with differential expression between persons with pulmonary NTM infections compared to other respiratory diseases (50). This suggests a possible role for peripheral blood transcriptional signatures as a biomarker of MAC lung disease. Such studies in TB indicate that signatures can predict progression to disease, distinguish between healthy and persons with TB disease, and associate with TB treatment status (51–57). Pursuit of peripheral blood transcriptional diagnostics for MACDZ may be an alternative to antigen-specific T-cell assays.

Limitations to our study population were inclusion of participants with cystic fibrosis, since they may have differences in immune responses. However, no differences in immune responses or RNA signatures were detected comparing those with and without cystic fibrosis. We were also unable to detect Mav-specific T-cell responses in MACDZ suggesting the need to examine individuals at earlier stages of infection, colonization, or disease.

In conclusion, this study provides a detailed characterization of immune responses in individuals with MAC disease. Peripheral signatures in MACDZ are characterized by impaired T-cell memory and hyperactive monocyte responses. These findings expand our understanding of the breadth of T-cell deficits associated with MAC disease extending to those without defined deficits. In addition, our data suggests a surprising parallel finding of enhanced innate responses which may be a critical new component of understanding MAC pathogenesis.

## Acknowledgements

We thank the sequencing core and flow cytometry core at La Jolla Institute for Immunology for their help in these studies.

## Supplemental material

### Methods

#### PBMC isolation and thawing

Venous blood was collected in heparin or EDTA containing blood bags or tubes and PBMCs were isolated by density gradient centrifugation using Ficoll-Hypaque (Amersham Biosciences) according to the manufacturer’s instructions. Cells were resuspended in FBS (Gemini Bio-Products) containing 10% DMSO (Sigma-Aldrich) and cryopreserved in liquid nitrogen.

Cryopreserved PBMC were quickly thawed by incubating each cryovial at 37°C for 2 min, and cells transferred to 9ml of cold medium (RPMI 1640 with L-glutamin and 25mM HEPES; Omega Scientific), supplemented with 5% human AB serum (GemCell), 1% penicillin streptomycin (Life Technologies), 1% glutamax (Life Technologies) and 20 U/ml benzonase nuclease (MilliporeSigma). Cells were centrifuged and resuspended in medium to determine cell concentration and viability using trypan blue and a hematocytometer.

#### Cell culture reagents, mycobacterial strains

Monocytes were cultured in Roswell Park Memorial Institute 1640 medium containing phenol red, HEPES and L-glutamine (RPMI 1640, Gibco) supplemented with fetal bovine serum (Atlas Biologicals) to a final concentration of 10% (RPMI-10) and recombinant human macrophage colony-stimulating factor (M-CSF, Peprotech) at 50 ng/mL. The *Mycobacterium avium* strain 104 (from the Cangelosi lab) was cultured in Middlebrook 7H9 media (BD Difco) supplemented with glycerol (Fisher; 4 mL/L), Middlebrook ADC Supplement (BD BBL, 100 mL/L) and Tween 80 (Fisher; 0.05% final) and grown to log-phase. Cultures were pelleted at 3,000 x g, washed twice in Sauton’s media, resuspended in Sauton’s media to OD ∼1.0 and aliquots were frozen at -80°C until monocyte infections. Freshly thawed *M. avium* 104 stocks were used to immediately infect monocyte cultures after obtaining the optical density to avoid heterogeneity between batches. The conversion of OD to CFU to achieve the desired multiplicity of infection (MOI) was determined by plating serial dilutions of a freshly frozen stock on Middlebrook 7H10 agar (BD BBL) for CFU enumeration.

#### Fluorospot assay

PBMCs were thawed and antigen-specific cellular responses were measured by IFN*γ*, IL- 5, and IL-17 Fluorospot assay with all antibodies and reagents from Mabtech (Nacka Strand, Sweden). Plates were coated overnight at 4°C with an antibody mixture of mouse anti-human IFN*γ* (Clone 1-D1K), IL-5 (TRFK5), and IL-17 (MT44.6). Briefly, 200,000 cells were plated in each well of the pre-coated Immobilon-FL PVDF 96-well plates (Mabtech), stimulated with the respective antigen (peptide pools at 1 μg/ml, Mav and Mtb whole cell lysates at 10μg/ml, PHA at 10μg/ml as a positive control and DMSO corresponding to the concentration present in the peptide pools). *M. avium* strain 104 whole cell lysate was prepared by heat-killing at 100°C for 25 minutes. Mtb whole cell lysate (Mtb lysate) from strain H37Rv was obtained from BEI Resources, NIAID, NIH (NR-14822). Fluorospot plates were incubated at 37°C in a humidified CO_2_ incubator for 20-24 hrs. All conditions were tested in triplicates. After incubation, cells were removed, plates were washed six times with 200 μl PBS/0.05% Tween 20 using an automated plate washer. After washing, 100μl of an antibody mixture containing anti-IFN*γ* (7-B6-1-FS-BAM), anti-IL-5 (5A10- WASP), and biotinylated anti-IL-17 (MT504) prepared in PBS with 0.1% BSA was added to each well and plates were incubated for 2 hrs at room temperature. The plates were again washed six times and incubated with diluted fluorophores (anti-BAM-490, anti-WASP-640, and anti-SA-550) for 1 hr at room temperature. After incubation, the plates were washed and incubated with a fluorescence enhancer for 15 min. Finally, the plates were blotted dry and spots were counted by computer-assisted image analysis (IRIS, Mabtech). The responses were considered positive if they met all three criteria (i) net spot forming cells per 10^6^ PBMC were *≥*20, (ii) stimulation index *≥*2, and (iii) p*≤*0.05 by Students’ t test or Poisson distribution test.

#### AIM assay

PBMCs were thawed and 1 million were added to a round-bottom 96-well plate and stimulated with the respective antigens (peptide pools, Mav and Mtb lysates, PHA and DMSO control). Plates were incubated at 37°C in a humidified CO_2_ incubator for 20-24 hrs. Plates were centrifuged and cells were resuspended in PBS with 10% (v/v) FBS and incubated at 4°C for 10 min. Cells were then stained with fixable viability dye eFluor506 (eBioscience) and an antibody mixture containing anti-human CD3-AF700 (UCHT1; ThermoFisher), CD4-APCeFluor780 (RPA- T4; eBioscience), CD8-BV650 (RPA-T8; BioLegend), CD14-V500 (M5E2; BD Bioscience), CD19- V500 (HIB19; BD Bioscience), CD25-PerCPCy5.5 (BC96; BioLegend), CD69-PECy7 (FN50; eBioscience), CD137-APC (4B4-1; BioLegend), CD154-FITC (TRAP1; BD Bioscience), OX40- BV421 (Ber-ACT35; BioLegend), and PD-L1-PE (29E.2A3; BioLegend) for 30 min at 4°C. Cells were washed, resuspended in PBS and acquired on a BD LSR II flow cytometer (BD Biosciences). Analysis to compare frequencies of activated CD4+ T cells was completed on FlowJo. The total number of CD4+ T cells expressing combinations of activation markers was determined with background values (as determined from the medium alone control) subtracted. The gating strategy is found in Figure S1A.

#### Intracellular cytokine staining

PBMCs were thawed and 1 million were added to a round-bottom 96-well plate and stimulated with the respective antigens (peptide pools, Mav and Mtb lysates, PHA and DMSO control). Anti-CD28 (1μg/ml, CD28.2; eBioscience) and anti-CD49d (1μg/ml, 9F10; BioLegend) were added to each well. Cells were incubated for 5 hours at 37°C. After 5 hrs, BFA (2.5μg/ml) and monensin (2.5μg/ml) were added and cells were incubated for an additional 7hrs at 37°C. Following the incubation cells were incubated with PBS with 10% (v/v) FBS at 4°C for 10min. They were then stained with fixable viability dye eFluor506 (eBiosciences) and an antibody mixture containing anti-human CD3-AF700 (UCHT1; ThermoFisher), CD4-APCeFluor780 (RPA- T4; eBioscience), CD8-BV650 (RPA-T8; BioLegend), CD14-V500 (M5E2; BD Bioscience), and CD19-V500 (HIB19; BD Bioscience) for 30 min at 4°C. After washing, cells were fixed using 4% paraformaldehyde and then permeabilized using saponin buffer (0.5% w/v saponin, 1% sodium acetate, 10% BSA in PBS). Cells were stained with CD154-PE (TRAP1; BD Bioscience), IFN*γ*- FITC (4S.B3; eBioscience), IL-2-PerCPeFluor710 (MQ1-17H12; eBioscience), and TNF*α*- eFluor450 (MAb11; eBioscience) in saponin buffer containing 10% FBS at room temperature for 20 min. Cells were washed, resuspended in PBS, and acquired on a BD LSR II flow cytometer (BD Bioscience). Combinations of cytokine production was determined using FlowJo and Boolean gating following the gating strategy in Figure S1B.

#### Flow cytometry for PBMC cell subsets

PBMCs were thawed and incubated in a round-bottom 96-well plate with PBS with 10% FBS for 10 min at 4°C. Cells were then stained with combinations of antibodies to determine PBMC cell subset frequencies. For CXCR3 and CCR6, stained cells were incubated at 37°C for 30min with anti-human CCR6-BV650 (G034E3; BioLegend) and CXCR3-APC (1C6-CXCR3; BD bioscience) before adding other antibodies. Cells were then stained with anti-human CCR4- PECy7 (1G1; BD Bioscience), CCR7-PerCPCy5.5 (G043H7, Biolegend), CD4-APCef780 (RPA- T4, eBiosciences), CD3-AF700 (UCHT1, BD Pharmigen), CD8a-BV650 (RPA-T8, Biolegend), CD19-PECy7 (HIB19, TONBO), CD14-APC (61D3), CD25-FITC (M-A251; BD Bioscience), CD45RA-eFluor450 (HI100, eBiosciences), CD127-PE (eBioRDR5, eBioscience), and eF506 live dead aqua dye (eBiosciences). The gating strategy is shown in Figure S1C.

In addition, other cells were stained with a mixture of the following antibodies: CD4- APCef780 (RPA-T4, eBiosciences), CD3-AF700 (UCHT1, BD Pharmigen), CD8a-BV650 (RPA- T8, Biolegend), CD19-PECy7 (HIB19, TONBO), CD14-APC (61D3), CCR7-PerCPCy5.5 (G043H7, Biolegend), CD56-PE (eBiosciences), CD25-FITC (M-A251, BD Pharmigen), CD45RA- eFluor450 (HI100, eBiosciences) and eF506 live dead aqua dye (eBiosciences, 65-0866-1) for 30 mins at 4°C. Cells were then washed twice and resuspended in 100 ul PBS for flow cytometric analysis on a BD LSRII flow cytometer (BD Bioscience). For this last panel we followed the previously described gating strategy (1).

#### Cell sorting and RNA purification

PBMCs were thawed and added at a density of 1×10^6^ cells per well to a round-bottom 96- well plate. They were stimulated for 24 hrs at 37°C with MTB300, Mav lysate, Mtb lysate, anti- CD28 (1μg/ml, CD28.2; eBioscience) together with anti-CD3 (1μg/ml pre-coated overnight at 4°C, UCHT1; BioLegend), as well as left unstimulated as a negative control. After 24 hrs, cells were resuspended in PBS with 10% (v/v) FBS and incubated at 4°C for 10 min. Cells were then stained with fixable viability dye eFluor506 (eBioscience) and an antibody mixture containing anti-human CD3-AF700 (UCHT1; ThermoFisher), CD4-APCeFluor780 (RPA-T4; eBioscience), CD8-V500 (RPA-T8; BD Horizon), CCR7-PerCPCy5.5 (G043H7, Biolegend) CD14-V500 (M5E2; BD Bioscience), CD19-V500 (HIB19; BD Bioscience), CD25-FITC (M-A251; BD Bioscience), CD45RA-eFluor450 (HI100, eBiosciences), CD137-APC (4B4-1; BioLegend), OX40-PECy7 (Ber- ACT35; BioLegend), and PD-L1-PE (29E.2A3; BioLegend) for 20 min at room temperature. Cells were washed and resuspended in PBS before being transferred into a 5 ml polypropylene FACS tube (BD Bioscience). PBMCs were sorted, based on forward and side scatter excluding debris and doublets (Figure S1D), on a FACSAria into TRIzol LS (Thermo Fisher). Acquisition files were analyzed using FlowJo. Total RNA was extracted from ∼100,000 cells in TRIzol LS using the miRNeasy Micro Kit (Qiagen) on a QIAcube (Qiagen). Total RNA was amplified according to Smart Seq protocol (2). cDNA was purified using AMPure XP beads. cDNA was used to prepare a standard barcoded sequencing library (Illumina). Samples were sequenced using an Illumina HiSeq2500 to obtain 50-bp single end reads. Samples that failed to be sequenced due to limited sample availability or failed the quality control were eliminated from further sequencing and analysis. The full protocol can be found at protocols.io (http://dx.doi.org/10.17504/protocols.io.bxr6pm9e).

#### RNA-sequencing data analysis

Paired-end reads that passed Illumina filters were filtered for reads aligning to tRNA, rRNA, adapter sequences, and spike-in controls. The reads were aligned to the GRCh38 reference genome and Gencode v27 annotations using STAR (v2.6.1) (3). DUST scores were calculated with PRINSEQ Lite v0.20.3 (4) and low-complexity reads (DUST > 4) were removed from BAM files. The alignment results were parsed via SAMtools (5) to generate SAM files. Read counts to each genomic feature were obtained with the featureCounts (v 1.6.5) (6) using the default option along with a minimum quality cut off (Phred > 10). After removing absent features (zero counts in all samples), the raw counts were imported into R v3.6.1 and genes with an average TPM < 1 were removed. R/Bioconductor package DESeq2 v.1.24.0 (7) was used to normalize raw counts. Variance stabilizing transformation was applied to normalized counts to obtain log2 gene expression values. Quality control was performed using boxplots and Principal component analysis (PCA), using the ‘prcomp’ function in R, on log_2_ expression values. Differentially expressed genes were identified using the DESeq2 Wald test, and p-values were adjusted for multiple test correction using the Benjamini Hochberg algorithm (8). Genes with adjusted p values < 0.05 and log2 fold change > 0.5 or < -0.5 were considered differentially expressed. Pathway enrichment analysis was performed using Enrichr (https://maayanlab.cloud/Enrichr/), and cell type enrichment was performed using DICE (9).

#### CD14+ monocyte isolation and M. avium infection

Peripheral blood mononuclear cells (PBMC) were isolated from selected individuals using Ficoll gradient separation, followed by washing, and cryopreservation. Cryopreserved PBMCs were thawed in batches of 8 donors (balanced by MAC subjects/healthy controls), (Day = 0) and viable cells, as assessed by Trypan Blue stain, were resuspended in RPMI/10 containing M-CSF (50 ng/mL) at 2 million cells per mL and rested overnight in non-TC treated dishes at 37_o_C/5% CO_2_. On day 1, CD14+ monocytes were enriched with negative selection using magnetic beads (Classical Monocyte Isolation Kit, Miltenyi Biotec) and then plated at 1 million cells per mL RPMI- 10 supplemented with M-CSF and again incubated at 37°C/5% CO_2_. The purity of the enriched CD14+ population was 60-80% as determined by flow cytometry. On day 2, cell cultures were stimulated either with *M. avium* 104 diluted in RPMI/10 to achieve an estimated MOI 5.0 or an equivalent volume of RPMI/10 media alone. After 6 hours, media was aspirated and cells were lysed in Trizol (Invitrogen) and lysates were transferred to cryotubes and stored at -80°C. RNA was isolated from lysates in batches by chloroform extraction and the application of the aqueous phase with 100% ethanol to miRNeasy mini columns, which were washed and eluted according to the manufacturer instructions (Qiagen). RNA quality was assessed by Agilent TapeStation to ensure RIN ≥ 8.0 and quantification was measured using Nanodrop (Thermo Scientific).

#### RNA sequencing and data processing for monocyte infection

Preparation of cDNA libraries, RNA sequencing and alignments using STAR2.6.0a. Sequences were quality-assessed with FastQC (v0.11.9 (10)) and filtered with AdapterRemoval (v2.3.2 (11)) to remove adapters and poor-quality sequences (score < 30, length < 15, ambiguous > 1). Sequences were aligned to the human genome (GRCh38 release 102) with STAR (v2.7.9a (3)) and quality-assessed with samtools flagstat (v1.7 (5)) and Picard (v2.42.2 (12)). Alignments were filtered with samtools to remove PCR duplicates, unmapped, non-primary, and poor-quality alignments (MAPQ < 30). High-quality alignments were then quantified in gene exons using Subread featureCounts (v2.0.1 (13)). Analysis and filtering steps were performed in R (v4.1.1 (14)). Counts were normalized for RNA composition using the trimmed mean of M-values normalization method and filtered to protein coding genes with at least 4% of libraries containing at least 1.5 count per million (CPM). Finally, counts were converted to log2 CPM using voom (15).

#### Differential gene expression, gene set enrichment analyses, and STRING network analysis

To identify genes with expression patterns that distinguished MAC and HC phenotypes according to the monocyte response to Mav infection, we selected a linear mixed effects model that incorporated an interaction term in addition to the main effects: Expression ∼ MACDZ + Mav + MACDZ:Mav +/- covariates with patient included as random effects and age, sex, and ethnicity included as covariates using R packages lme4 (16). Inclusion of age, sex, or ethnicity as covariates in the model did not improve the fit (median sigma changes 0.0001 for age, 0.00007 for sex, and 0.000003 for ethnicity, Figure S4A-C). Furthermore, except for ethnicity, no clustering was detected on PCA plots of these covariates (Figure S5). Differentially expressed genes were assessed at an FDR<0.05, and significant genes were further assessed in a MACDZ:Mav pairwise contrasts model including MACDZ within media or Mav-infected and Mav infection within HC or MACDZ.

We also explored whether to include the samples from cystic fibrosis (CF) subjects due to imbalance of this variable in the cases and controls (6 vs 0, respectively). Using pairwise comparisons of MACDZ subjects with and without CF, we did not discover any DEGs in the media or Mav condition. In addition, there was no difference in CF vs no-CF clustering on a PCA plot (Figure S6D) or improved model fit with exclusion of CF samples (Figure S5D&E). However, removal of CF samples lowered the numbered of DEGs substantially (227 vs 45 at FDR <0.05) likely due to reduced power. Without evidence of confounding by the CF samples, we proceeded with further analyses with inclusion of the CF samples.

To understand biologic connectivity between significant genes, we used STRING v11 network analysis (17) of Mav-dependent DEGs as defined by the interaction term (2 genes) or both MACDZ and Mav infection (87 genes), as well as Mav-independent DEGs as defined by MACDZ alone (138 genes). We identified one major cluster for Mav-dependent DEGs (31 out of 89 genes) and one for Mav-independent DEGs (28 out of 138 genes. DEGs were also assessed for enrichment against Gene Ontology (GO), Hallmark and KEGG gene sets using Fisher’s exact test in Enricher (18). Gene set enrichment analysis (GSEA) was performed using the Molecular Signatures Database (MSigDB v7.2 (19)) Hallmark and Gene Ontology (GO) collections. Fast gene set enrichment analysis (FGSEA (20)) was used to compare fold changes of all genes in MACDZ:Mav pairwise contrasts as described above. Leading-edge genes in significant GSEA results (FDR < 0.1) were compared between MAC and HC to identify significant pathways.

## Supplemental Figures and Legends

**Figure S1.**
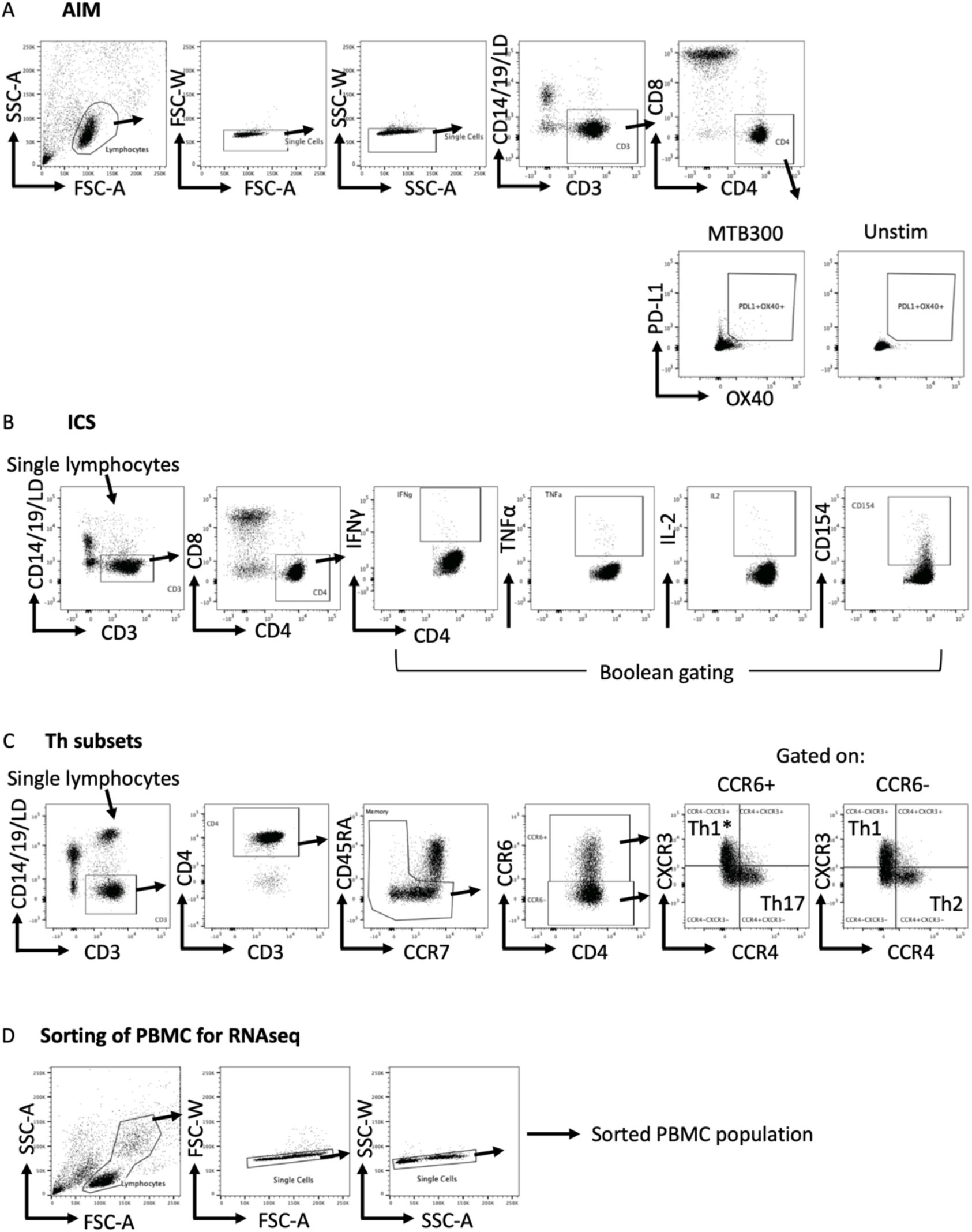
Gating strategy for flow cytometry experiments. A) AIM assay, B) ICS assay, C) Th subset determination, D) Sorting strategy. A-C) Live/Dead stain (LD), B,C) The single lympohocyte population was gated based on forward and side-scatter parameters

**Figure S2.**
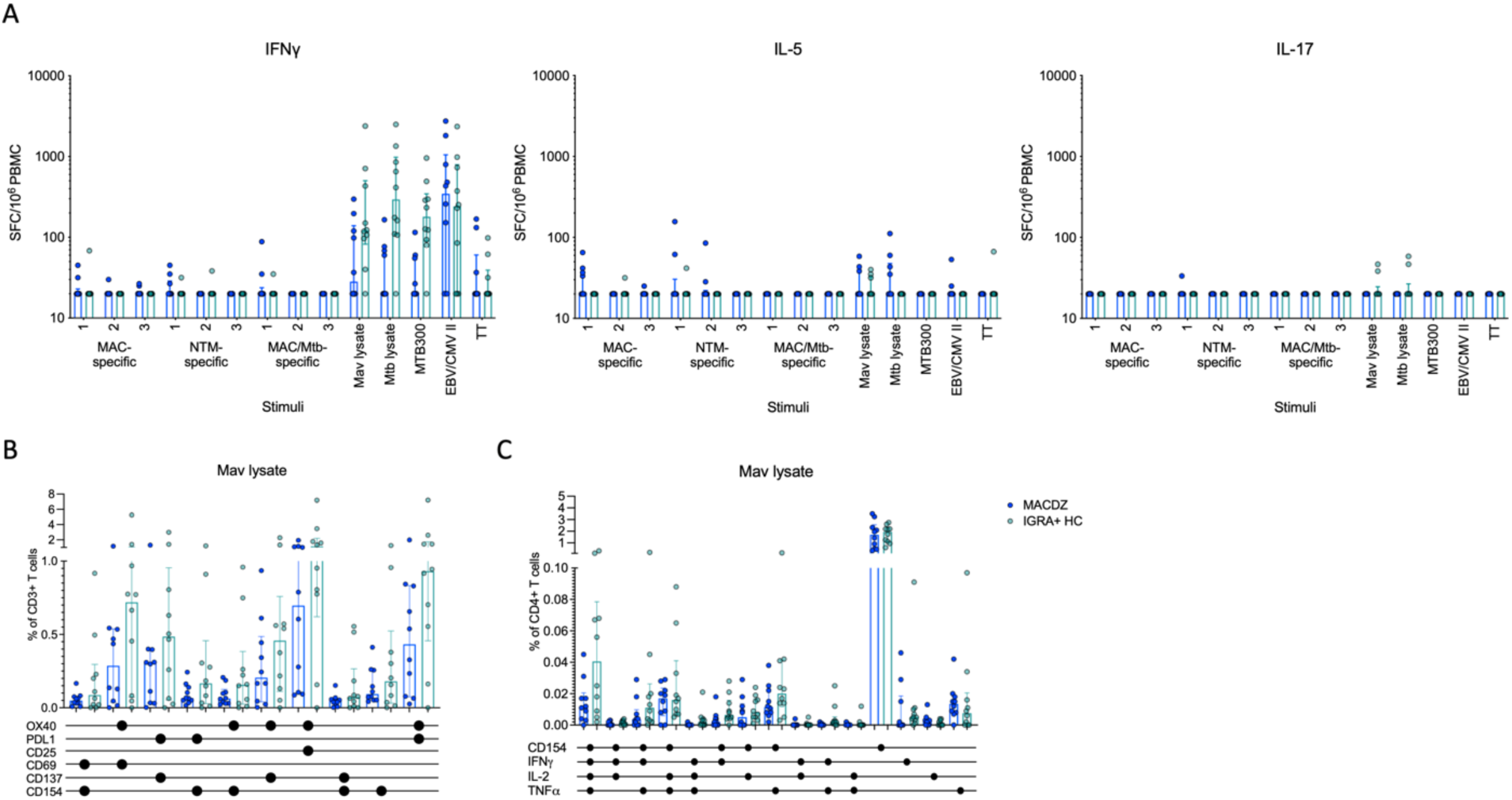
Individuals with MAC disease have infrequent Mav- or mycobacteria-specific responses. A) IFN*γ*, IL-5 or IL-17-specific magnitude of response against peptide pools, Mav and Mtb lysates as SFC per 10^6^ cultured PBMC as determined by Fluorospot. Each point represents one participant (MACDZ, n=10 in blue; IGRA+HC, n=10 in teal), median ± interquartile range is shown. B) Frequency of indicated combinations of activation markers in CD4 T-cells in response to Mav lysate shown as % of CD3+ T-cells. Each point represents one participant (MACDZ, n=10; IGRA+HC, n=10), median ± interquartile range is shown. Two-tailed Mann-Whitney test is >0.05 for all stimuli comparing MACDZ vs. IGRA+HC. CD4 T-cells were gated as CD3+CD4+CD8-CD19-CD14- in the live singlet gate of PBMC. C) Frequency of indicated combinations of CD154, IFN*γ*, IL-2 and TNF*α* in CD4 T-cells in response to Mav lysate. Each point represents one participant (MACDZ, n=10; IGRA+HC, n=10), median ± interquartile range is shown. Combinations of cytokines were determined by Boolean gating following the gating strategy in Figure S1.

**Figure S3.**
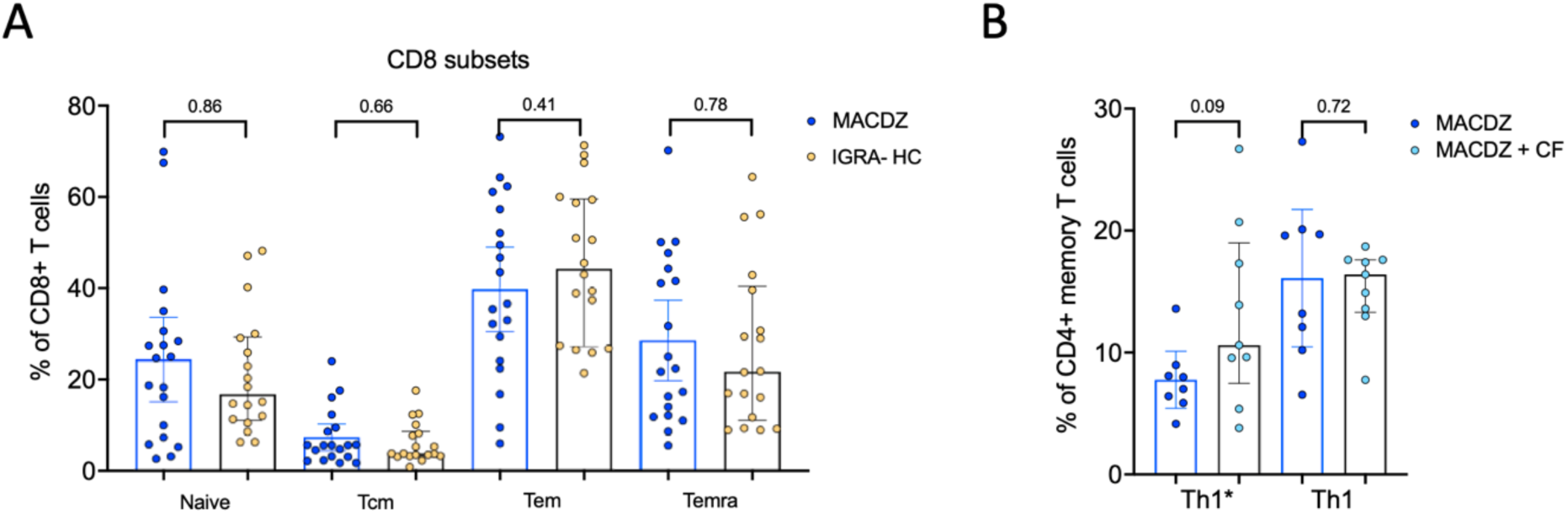
Individuals with MAC disease have similar frequencies of CD8 memory populations and no difference if they have cystic fibrosis of specific cell subsets. A) Frequency of CD8 memory populations based on CD45RA and CCR7 expression divided in naïve, effector memory (Tem), central memory (Tcm) and Temra populations. Each point represents one participant (MACDZ, n=19 in blue, IGRA-HC, n=18 in yellow), median ± interquartile range is shown. Two-tailed Mann-Whitney test. B) Frequency of specific Th subsets (Th1* and Th1) in MACDZ individuals with (MACDZ+CF, n=9 in light blue) or without cystic fibrosis (MACDZ, n=8, in dark blue). Each point represents one participant, median ± interquartile range is shown. Two-tailed Mann-Whitney test.

**Figure S4.**
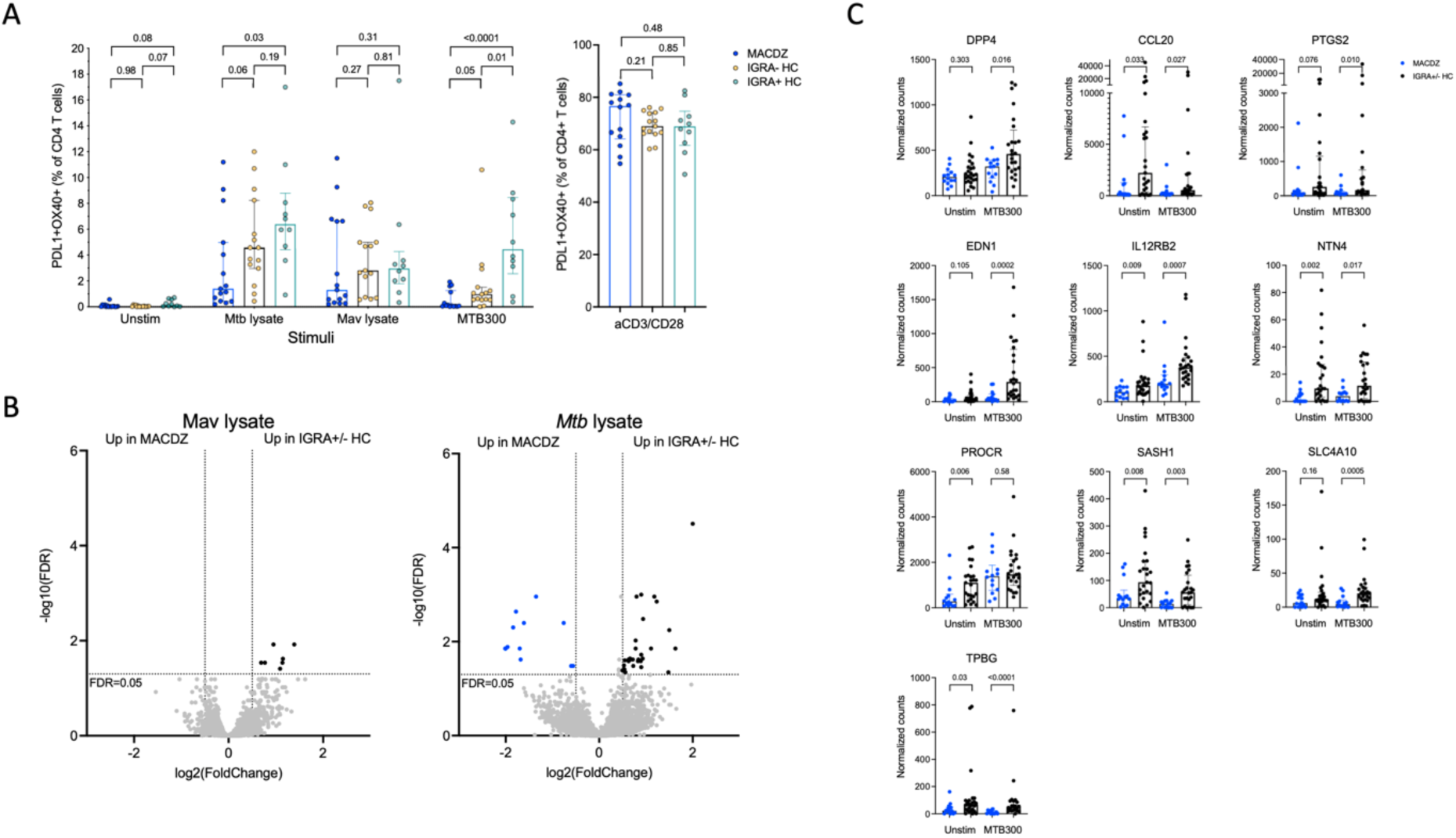
Activation induced marker upregulation in RNAseq cohorts, and differentially expressed genes in response to Mav and Mtb lysates. A) Frequency of PDL1+OX40+ CD4 T-cells in response to MTB300, Mav and Mtb lysates, and anti- CD3/CD28 stimulation. Each point represents one participant (MACDZ, n=15; IGRA-HC, n=10, and IGRA+HC, n=10), median ± interquartile range is shown. Two-tailed Mann-Whitney test. CD4 T-cells were gated as CD3+CD4+CD8-CD19-CD14- in the live singlet gate of PBMC. B) Volcano plots showing differentially expressed genes in Mav and Mtb lysate stimulated samples comparing individuals with MAC disease (upregulated genes shown in blue) to IGRA+/-HC individuals (upregulated genes shown in black). Adjusted p-value <0.05 and log2 fold change >0.5 or <-0.5, Benjamini Hochberg corrected DESeq2 Wald test. C) Th1* signature gene expression in the unstimulated or MTB300 stimulated MACDZ and IGRA+/-HC samples. Each point represents one participant, median ± interquartile range is shown. Two-tailed Mann-Whitney test.

**Figure S5.**
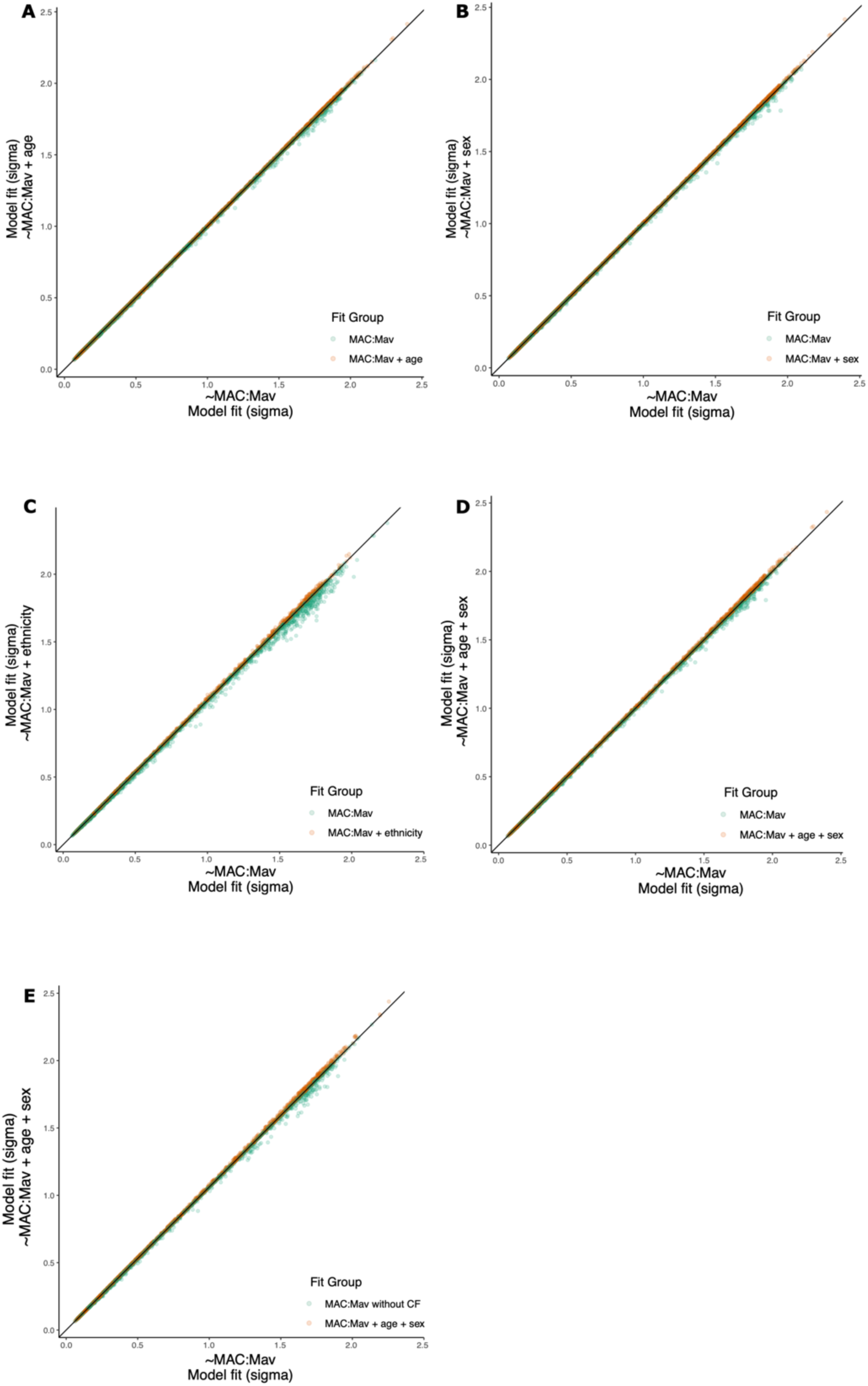
Sigma plots from linear model comparing RNASeq transcriptional profiles in MACDZ and HC subjects. Expression profiles were compared between MACDZ and HC subjects with and without Mav infection using a linear model that incorporated an interaction term in addition to the main effects: Expression ∼ phenotype + stimulation + phenotype:stimulation and Expression ∼ MACDZ + Mav + MACDZ:Mav +/- covariates with patient included as random effects and age, sex, and ethnicity included as covariates using R packages lme4. (A-C) Sigma plots indicate that inclusion of age **(A)**, sex **(B)**, or ethnicity **(C)** as covariates in the model did not improve the fit (median sigma changes 0.0001 for age, 0.00007 for sex, and 0.000003 for ethnicity. To assess the effect of CF, we examined sigma plots of a model including age and sex as covariates with **(D)** and without **(E)** CF samples. The model fit did not improve with including age, sex, or ethnicity as covariates or with the exclusion of CF samples.

**Figure S6.**
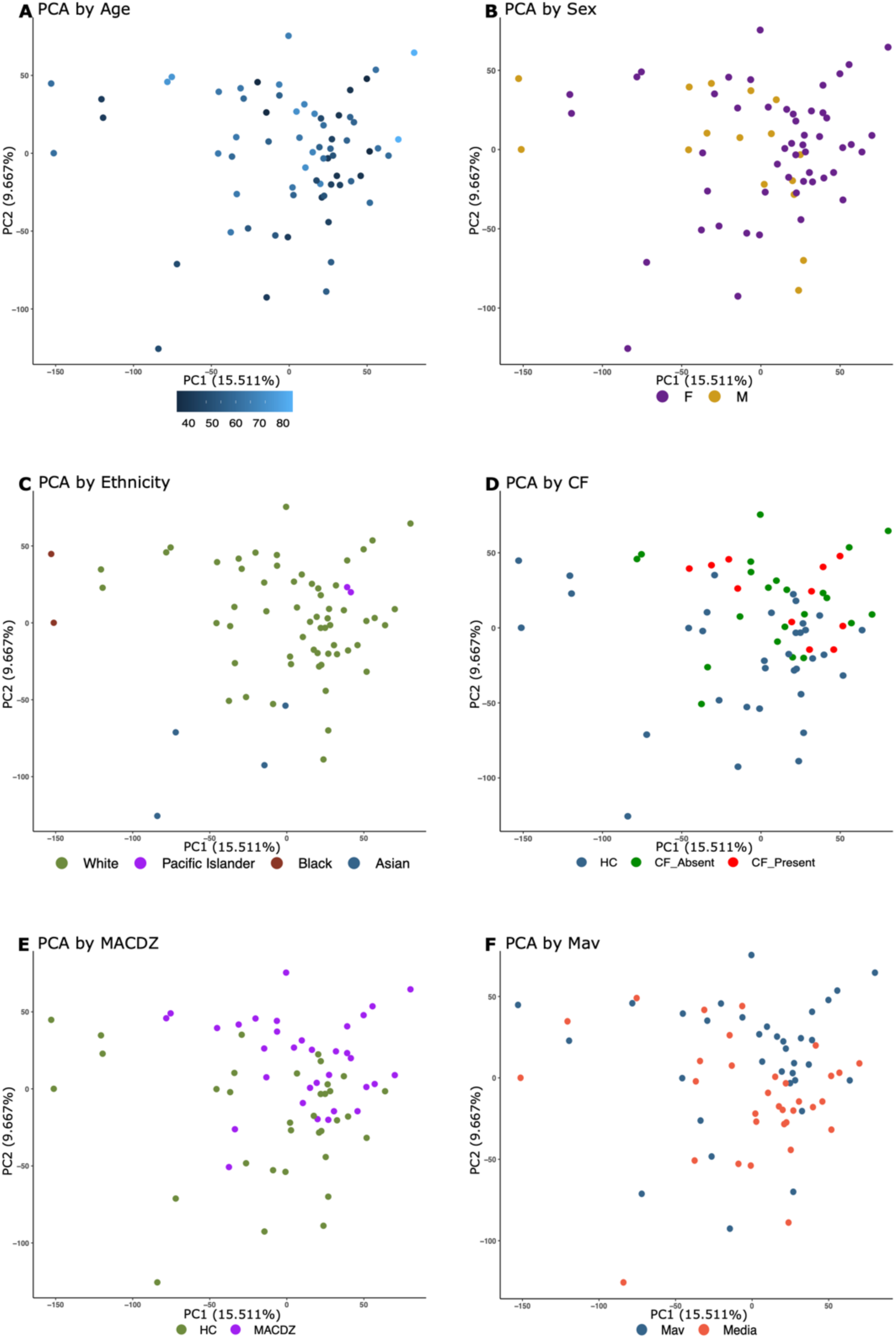
Principal component analysis (PCA) of RNASeq profiles of monocyte response to *Mycobacterium avium* infection in MACDZ versus HC subjects. Principal component analysis (PCA) plots of monocyte RNASeq transcriptional profiles from MACDZ subjects (N=17) or HC (N=17) with media only condition or after infection with *M. avium* (MOI=5) for 6 hours. PCA colored by age **(A)**, sex **(B)**, ethnicity **(C)**, cystic fibrosis **(D),** MACDZ vs HC **(E)**, or media vs M. avium condition **(F)**.

**Figure S7.**
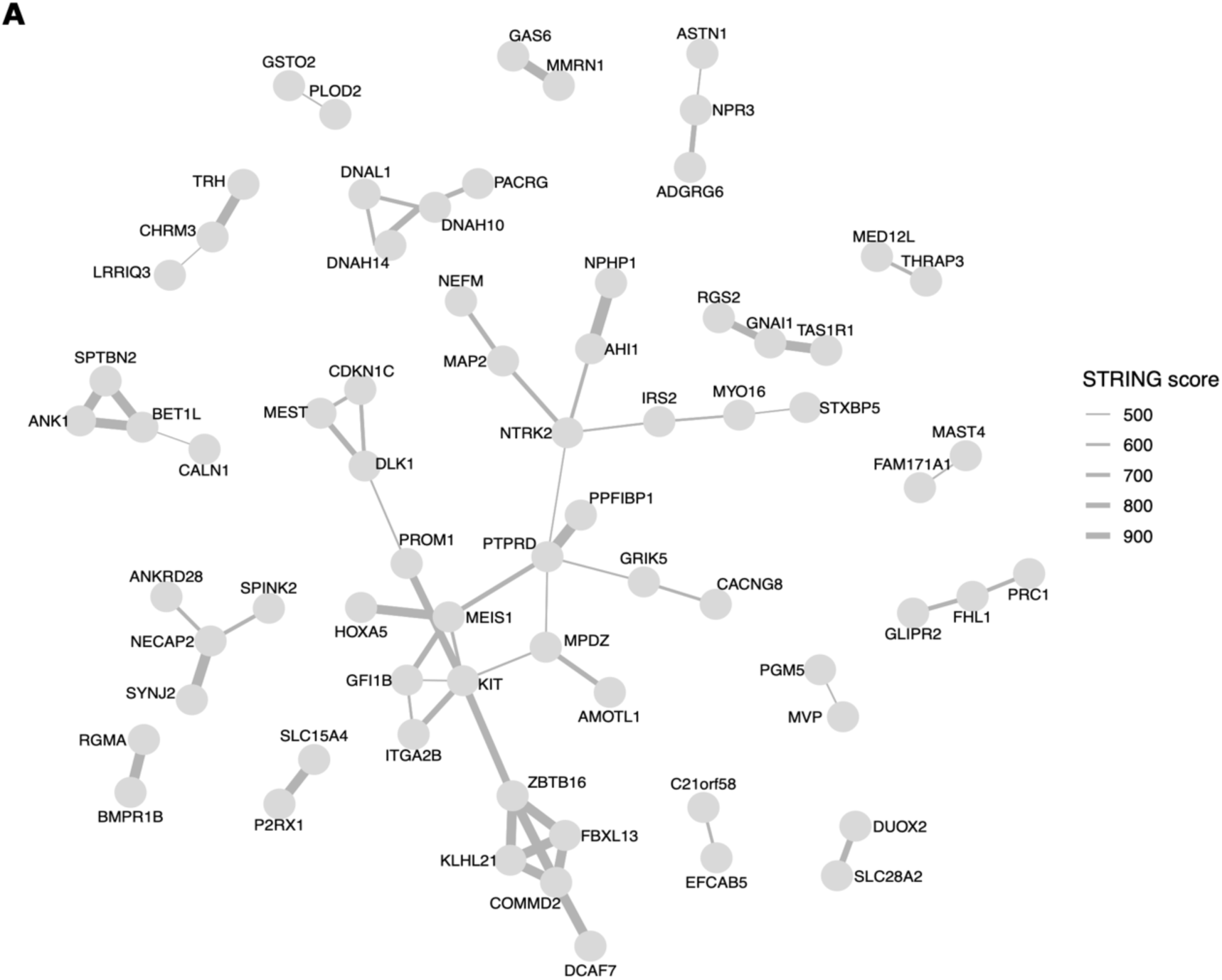
STRING network of media condition DEGs from monocyte transcriptional profile in MACDZ versus HC subjects. STRING network analysis of 138 Mav-independent monocyte DEGs which differentiate MACDZ vs HC subjects. Circles depict genes. Grey lines depict annotated connection between 2 genes in the STRING database with line thickness proportionate to the score.

**Figure S8.**
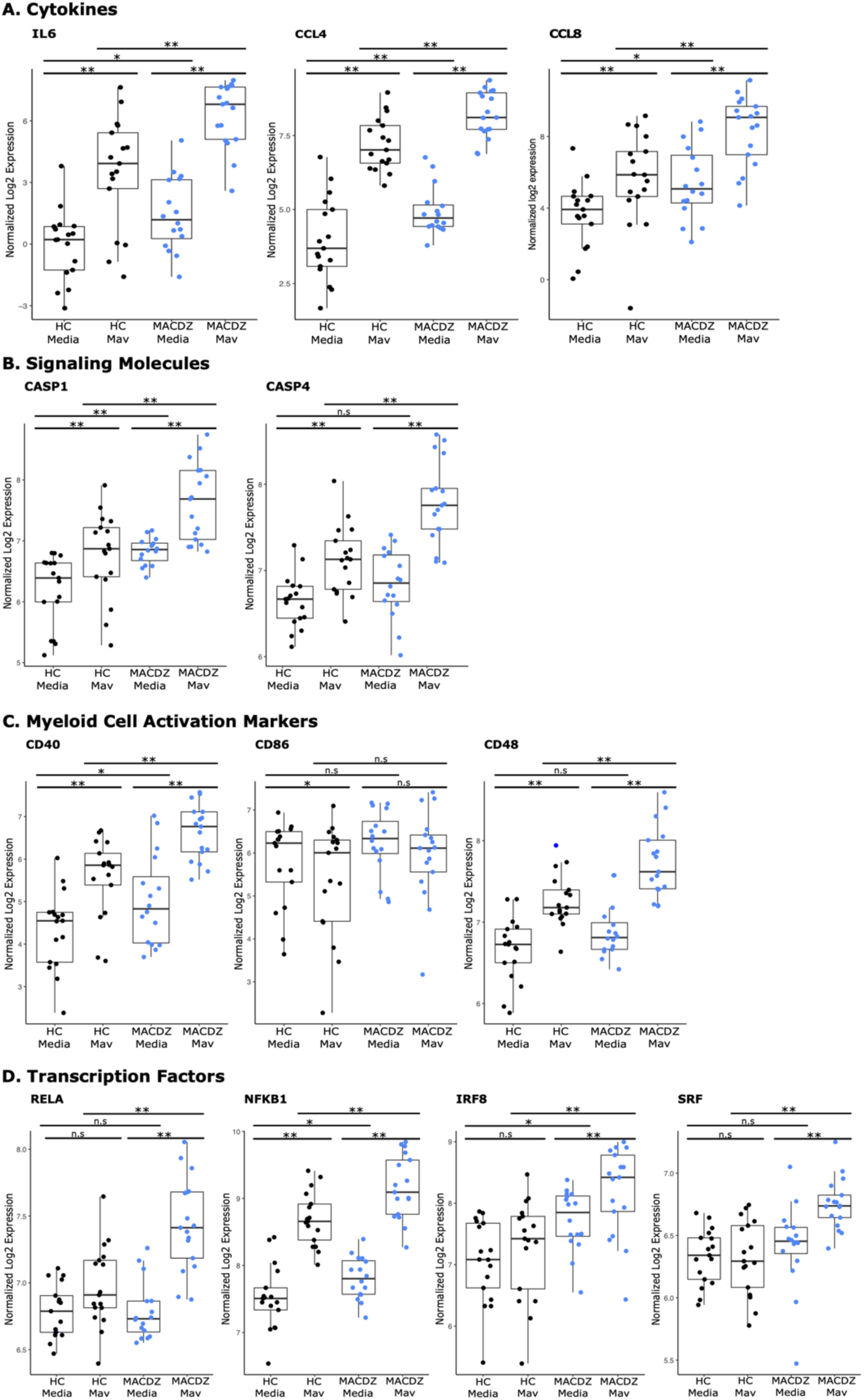
DEG boxplots from monocyte transcriptional response to *Mycobacterium avium* infection in MACDZ versus HC subjects. Boxplots depicting voom normalized log2 mRNA expression values in MACDZ versus HC subjects with and without Mav infection. FDR values depict comparison of MACDZ vs HC media expression, MACDZ vs HC Mav expression, and Mav vs media for MACDZ or HC subjects (FDR*≤*0.1 shown by •, FDR*≤*0.05 shown by *, and FDR*≤*0.01 shown by **). Median and interquartile range depicted. Values were higher in the MACDZ subjects compared to HC for the media and/or Mav condition. Genes are expanded number and identical dataset as Fig. 4D-F.

## Supplemental Tables

**Table S1. List of peptide sequences included in the peptide library**

**Table S2. Differentially expressed genes identified in the RNAseq analysis of PBMCs**

**Table S3. Differentially expressed genes from linear model with interaction term comparing MACDZ versus HC monocyte transcriptional profiles with media and *M. avium* infection conditions.** The table is split into three tabs for differentially expressed genes in MACDZ, Mav-dependent and interaction term groups, respectively.

**Table S4. Gene set enrichment analysis with Hallmark mSigDB terms of MAC versus healthy control monocyte transcriptional profiles with media and *M. avium* infection conditions.** The table is split into two tabs for pathways enriched in MACDZ vs. HC within *M. avium* infection and MACDZ vs. HC within media condition.

**Table S5. Hypergeometric mean pathway enrichment analysis of 138 Mav- independent and 89 Mav-dependent DEGs of monocyte transcriptional profiles of MACDZ versus HC subjects with media and *M. avium* infection conditions.** The table has separate tables for Hallmark and KEGG gene sets both for Mav-independent and Mav-dependent DEGs.

